# PIN-FORMED1 polarity in the shoot is insensitive to the polarity of neighbouring cells

**DOI:** 10.1101/2021.09.27.462050

**Authors:** Abdul Kareem, Neha Bhatia, Carolyn Ohno, Marcus G. Heisler

## Abstract

In plants, epidermal cells are planar-polarized along an axis marked by the asymmetric localization patterns of several proteins including PIN-FORMED1 (PIN1), which facilitates the directional efflux of the plant hormone auxin to pattern phyllotaxis (Heisler et al., 2010; Mansfield et al., 2018; Reinhardt et al., 2003). While PIN1 polarity is known to be regulated non-cell autonomously via the MONOPTEROS (MP) transcription factor, how this occurs has not been determined (Bhatia et al., 2016). Here we use mosaic expression of the serine threonine kinase PINOID (PID) to test whether PIN1 polarizes according to the polarity of neighbouring cells. Our findings reveal that PIN1 is insensitive to the polarity of PIN1 in neighbouring cells arguing against auxin flux or extracellular auxin concentrations acting as a polarity cue, in contrast to previous model proposals (Abley et al., 2016; Stoma et al., 2008).

## Introduction

Unlike animals, plants typically generate new lateral organs throughout post-embryonic development, often in a periodic fashion. The spatial organisation of these organs, also called phyllotaxis, has been a focus of intense interest for biologists, mathematicians, and physicists alike for many years (Jean and Barabé, 1998). Plant organogenesis is triggered by the plant hormone auxin and in the shoot, auxin is concentrated at organ initiation sites through a polar auxin transport system that depends on the membrane-bound auxin efflux carrier PIN-FORMED1 (PIN1) (Kuhlemeier and Reinhardt, 2001; Okada et al., 1991; Reinhardt et al., 2000). PIN1 directs auxin flux according to its asymmetric or polar localisation and in meristem epidermal cells at a supra-cellular scale, PIN1 polarity forms convergence patterns oriented toward both organ initiation sites as well as the shoot apex (Galvan-Ampudia et al., 2020; Heisler et al., 2005b; Mansfield et al., 2018; Reinhardt et al., 2003).

Given that polarity convergence patterns dictate where auxin accumulates and therefore where new organs form, in order to understand plant phyllotaxis it is necessary to understand how such polarity patterns are generated. Through computational modelling, a number of studies have shown that feedback between auxin and the polarity of its transporter protein PIN1 can account for PIN1 convergence patterns (Abley et al., 2016; Cieslak et al., 2015; Hartmann et al., 2019; Heisler et al., 2010; Heisler and Jonsson, 2006; Smith et al., 2006; Stoma et al., 2008). These models differ in the way auxin is proposed to provide polarity information. For instance “up the gradient” models propose that cells polarize their PIN1 towards neighbouring cells in proportion to the relative internal auxin concentrations (Jonsson et al., 2006; Smith et al., 2006) with mechanical signals acting to mediate cell-cell communication (Heisler et al., 2010). Another set of models propose auxin regulates PIN1 polarity via its net flux across the plasma membrane or as an extracellular coupling factor that regulates intra-cellular polarity partitioning (Abley et al., 2013; Abley et al., 2016; Stoma et al., 2008). A central prediction of the latter proposals is that the polarity of PIN1 is sensitive to the polarity of PIN1 in neighbouring cells, assuming PIN1 in these cells is transporting auxin (Abley et al., 2013; Abley et al., 2016).

Although the above-mentioned classes of models are relatively well established in the literature, few studies have tested their assumptions experimentally. In an effort to test the “up the gradient” model Bhatia et al., engineered differences in auxin signalling between neighbouring cells by inducing clonal expression of the auxin response factor MONOPTEROS (MP) in *mp* mutants. These experiments demonstrated that local differences in auxin signalling between cells can indeed act as a polarity cue, as “up the gradient” models would predict (Bhatia et al., 2016). However the observed reorientation of PIN1 polarity towards MP expressing cells can also be explained by flux-based and auxin indirect coupling models if local MP expression induces expression of auxin influx carriers and auxin degradation or conjugation enzymes, such that there is net auxin influx (Abley et al., 2016). Furthermore, it is possible to envision a scenario in which mechanics plays a role to orient a polarity axis while flux determines PIN1 mediated efflux direction (Marconi et al., 2021). Here we test indirect coupling models and models based on auxin flux by assessing the sensitivity of PIN1 polarity to the polarity of PIN1 in neighbouring cells in the *Arabidopsis* shoot using mosaic expression of the serine threonine kinase PINOID (PID).

## Results and Discussion

The PID serine threonine kinase is known to directly phosphorylate PIN1 (Christensen et al., 2000; Michniewicz et al., 2007) and its activity is both necessary and sufficient to promote apical polarization of PIN1 in the root and embryo (Friml et al., 2004). In the shoot PID also promotes a convergent or apical polarity since in *pid* mutants, PIN1 is predominantly polarized in a divergent pattern away from the apex and toward the root (Friml et al., 2004) (Fig. 1a). In addition to promoting apical polarization, PID activity also enables PIN1 localization to respond to mechanical signals (Friml et al., 2004; Heisler et al., 2010), such as those involved in organ formation (Hamant et al., 2008). We note that besides regulating PIN1 polarity, PID has been implicated in activating PIN1-mediated auxin efflux, as assessed in heterologous assays using Xenopus oocytes (Zourelidou et al., 2014). However genetic data strongly indicate that *in planta*, PIN1 retains efflux activity in the absence of PID, possibly due to the redundant activities of PID2, WAG1 and WAG2 (Cheng et al., 2008; Reinhardt et al., 2003).

**Figure 1.**
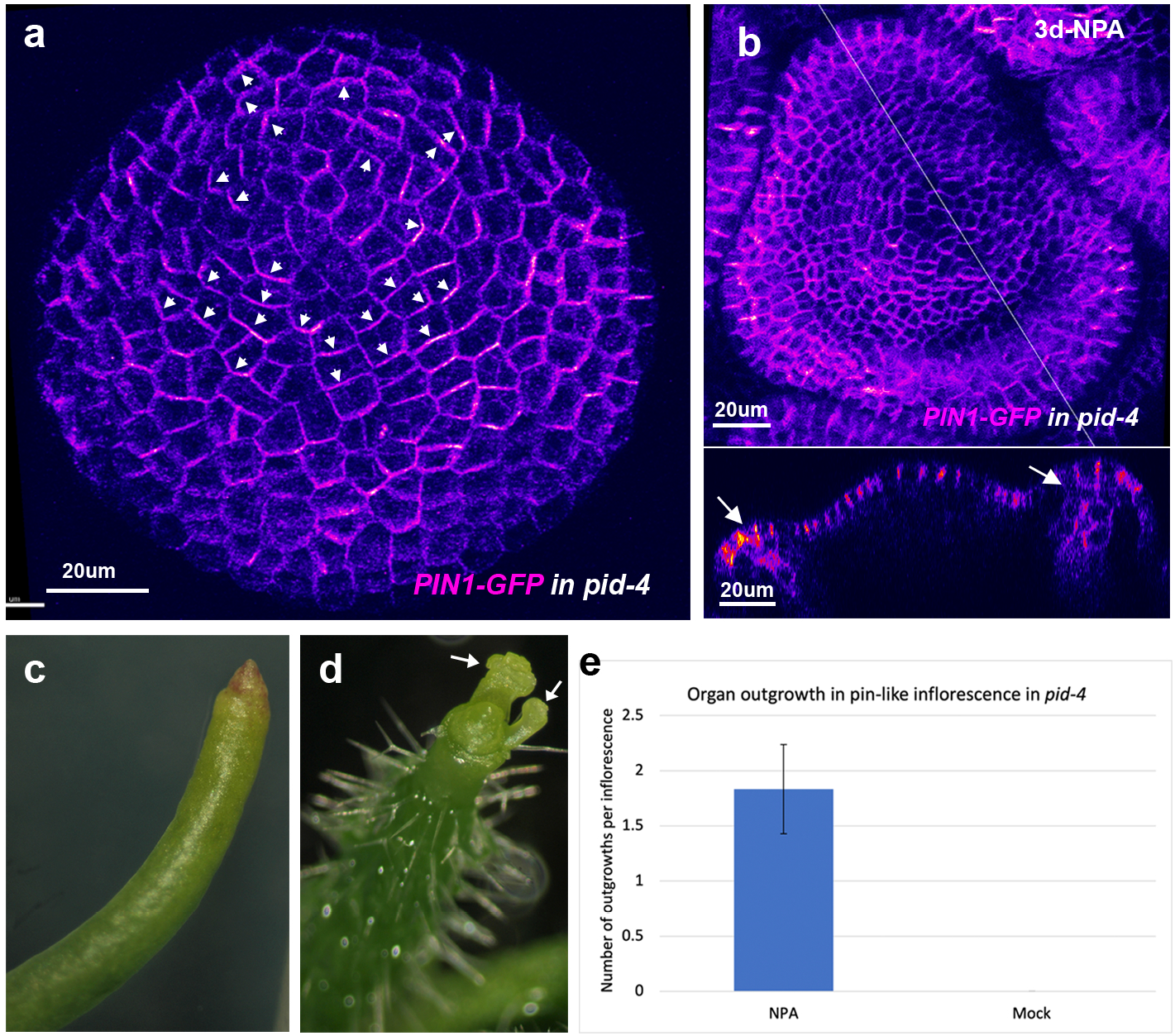
Basally localized PIN1 in pid mutant meristem transports auxin basally. **a**, *PIN1::PIN1-GFP* expression (magenta) in *pid-4* mutant inflorescence meristem. Basal PIN1 polarity away from meristem centre marked by arrowhead. **b**, *pid-4 PIN1-GFP* inflorescence meristem forming floral organ tissue 3 days after 10μM NPA treatment. Lower image shows longitudinal optical section of the meristem corresponding to line in top image. Arrows indicate organ primordia. **c**, Mock treated *pid-4 PIN1-GFP* inflorescence. **d**, *pid-4 PIN1-GFP* apex treated with 10μM NPA for 7 days. Arrows indicate floral primordia. **e**, Graph showing number of organ primordia produced from the *pid-4* mutant apex after NPA and mock treatments (n=12). Error bar represents standard error of mean.

Auxin signalling is relatively low in the *pid* mutant apex (Galvan-Ampudia et al., 2020), as might be expected if PIN1 transports auxin basally due to its basal polarity. To further investigate this possibility, we grew *pid-4* plants on 10 uM n-naphthylphthalamic acid (NPA) to see whether reductions in auxin transport could alter the flowerless *pid-4* phenotype and found that flower formation was partially rescued (Fig. 1b-e). While 75% of NPA treated pin-like inflorescences produced an average of 1.8 flowers per inflorescence (n=−12), mock treated inflorescences did not make any flowers (n=12) (Fig. 1c). These data support the proposal that in *pid-4* mutants, basally localized PIN1 transports auxin basally, thereby depleting auxin from the meristem and causing a flowerless phenotype.

Given PIN1 is active in *pid* mutants and that in the wild type, PID is present throughout the shoot epidermis, albeit at higher levels in boundary regions (Fig. 2a), we sought to restore local PID expression in *pid* mutants to enable proper polarization of PIN1 and allow us to assess whether PIN1 polarizes according to the prevailing pre-existing polarity of the tissue. Using an inducible system for Cre-lox recombinase mediated recombination, we generated epidermal clones of cells constitutively expressing PID fused to two copies of VENUS (PID-2V) in *pid* mutant SAMs. We found epidermal PID expression to be sufficient for rescuing organ formation since when large sectors expressing PID-2V were generated, organs formed that were marked by PID-2V expression (Fig. S1a-d). Closer examination also revealed local PID-2V expression associated with the formation of polarity convergence patterns at the SAM periphery (Fig. 2b-d, Movie-1).

**Figure 2.**
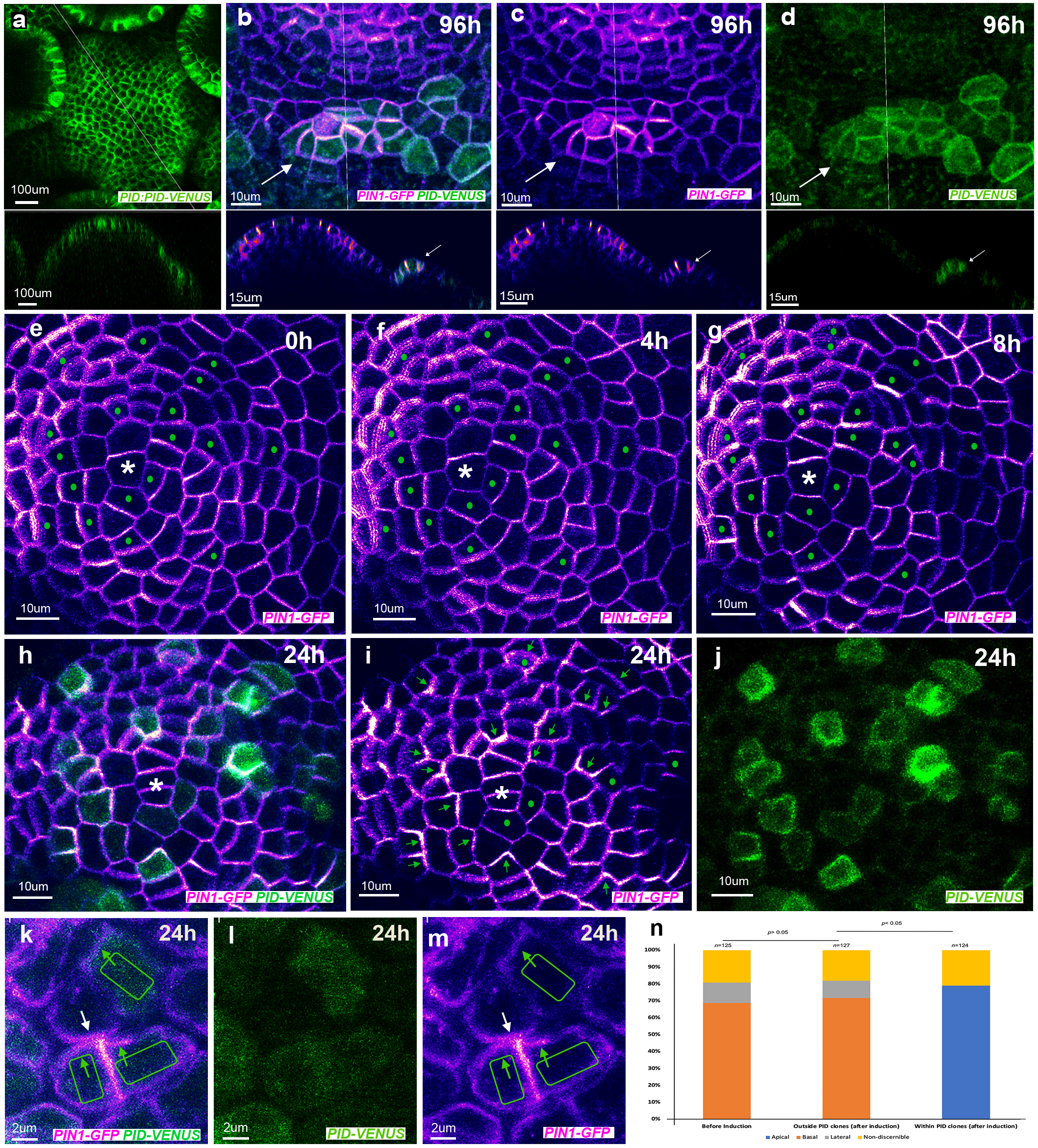
PID promotes changes in PIN1 polarity irrespective of initial or neighboring cell polarities. **a**, Expression pattern of *PID::PID-2V* (green) in wild type inflorescence meristem. Lower image represents longitudinal optical section corresponding to line in the upper image. **b-d,** Organ primordium (arrow) associated PIN1 convergence (magenta) co-localized with clonal PID expression (green) 96h after clone induction. Lower images in **b-d** represent longitudinal optical sections corresponding to line in upper images. PID-2V and PIN1-GFP (**b**), PIN1-GFP (**c**) and PID-2V (**d**). **e-j,** Time-lapse images of the *pid-4* meristem before (**e**) and after (**f-j**) induction of *PID-2V* (green) expressing clones showing changes in *PIN1-GFP* (magenta) polarity. The time interval of imaging: 0h (**e**), 4h (**f**), 8h (**g**) and 24h (**h-j**). Note apical shift in PIN1-GFP polarity (green arrows in **i**) in cells expressing PID-2V (green dots in **e-g**). Green dots in (**i**) mark cells with unclear polarity. Asterisk marks middle of the meristem. **k-m,** Magnified high resolution images of plasmolysed *pid-4* mutant meristem showing opposing PIN1-GFP (magenta) polarities (green and white arrows in (**k**) and (**m**)) in adjacent cells due to differential PID-2V expression (green) 24h after induction. PID-2V and PIN1-GFP (**k**), PID-2V (**l**) and PIN1-GFP (**m**). **n**, Quantification of the shift in PIN1 polarity in *pid-4* meristem after the induction of PID-2V (n=124).

Next, we examined PIN1 signal within membrane-localized PID-2V marked cells over time, from when PID-2V signal first became detectible 6-8 hrs after clone induction. We found a distinct basal to apical shift in PIN1 polarity in PID-2V marked cells, even when such cells were surrounded by PID negative cells that were polarized basally (Fig. 2e-j, Fig. 2n, Fig. S1e-r, Movie-2). We also found that PID positive clones did not influence the polarity of PIN1 in mutant neighbouring cells, which remained basal (Fig. 2k-m, Fig. S1s-u).

All together these results indicate that in the plant epidermis, where PID is normally expressed, the polarity of PIN1 in an apical or convergent orientation, which is critical for phyllotaxis, does not necessarily depend on the polarity of the auxin efflux carrier PIN1 in neighbouring cells. These findings contrast to predictions based on flux-based or indirect coupling models using auxin acting as the coupling agent (Abley et al., 2013; Abley et al., 2016; Stoma et al., 2008). Instead, our data indicate auxin likely influences cell polarity via additional intercellular signals, such as mechanical stress.

## Materials and Methods

### Plant material and growth conditions

The *pid-4* mutant *(Ler ecotype)* allele was previously described(Bennett et al., 1995). Transgenic lines used in this study include *pid-4* mutant transformed with *pPIN1::PIN1-GFP* reporter and *pid-4 PIN1::PIN1-GFP* transformed with *pML1::CRE-GR+pUBQ10::lox-GUS-lox-PID-2XVENUS.*

Seeds were germinated and grown on growth medium (GM) containing 1X Murashige and Skoog (MS) basal salt mixture (Sigma M5524), 1% sucrose, 0.5% MES 2-(MN-morpholino)-ethane sulfonic acid (Sigma M2933), 0.8 % Bacto Agar (BD) and 1% MS vitamins (Sigma M3900). pH was adjusted to 5.7 with 1M KOH. Plants were grown at 22°C under continuous light. For imaging of wild type inflorescence meristems, plants were grown on soil at 18°C in short day conditions (16h/8h).

### Construction of reporters and transgenic plants

For mosaic analyses using the CRE/Lox system, stable transgenic lines harbouring a template for sectoring (UBQ10p::lox spacer lox::PID-2XVENUS) and a dexamethasone-inducible CRE Recombinase (ML1p::CRE-GR) were used. To generate UBQ10p::lox spacer lox::PID-2XVENUS, first a 4.6 kb SfiI-BamHI fragment from UBQ10p::lox spacer lox::MP-VENUS(Bhatia et al., 2016) was cloned upstream of a 9X alanine linker followed by 2 tandem copies of VENUS and OCS terminator. PID cDNA sequence was amplified with primer set 121 and 122 and then cloned BamHI fragment as a translational fusion to 2XVENUS to create UBQ10p::lox spacer lox::PID-2XVENUS. CRE-GR(Bhatia et al., 2016) was cloned as a BglII fragment downstream 3.38 kb of ATML1 (At4g21750) 5’-regulatory sequences amplified with primer set 0708H3f and 0708Br to generate ML1p::CRE-GR. Finally, the UBQ10p::lox spacer lox::PID-2XVENUS, and ML1p::CRE-GR were combined in T-DNA vector BGW (Karimi et al., 2002) by Gateway technology (Invitrogen). The constructs were further transformed into agrobacterium strain C58C1 by electroporation and then finally transformed into Arabidopsis by floral dipping.

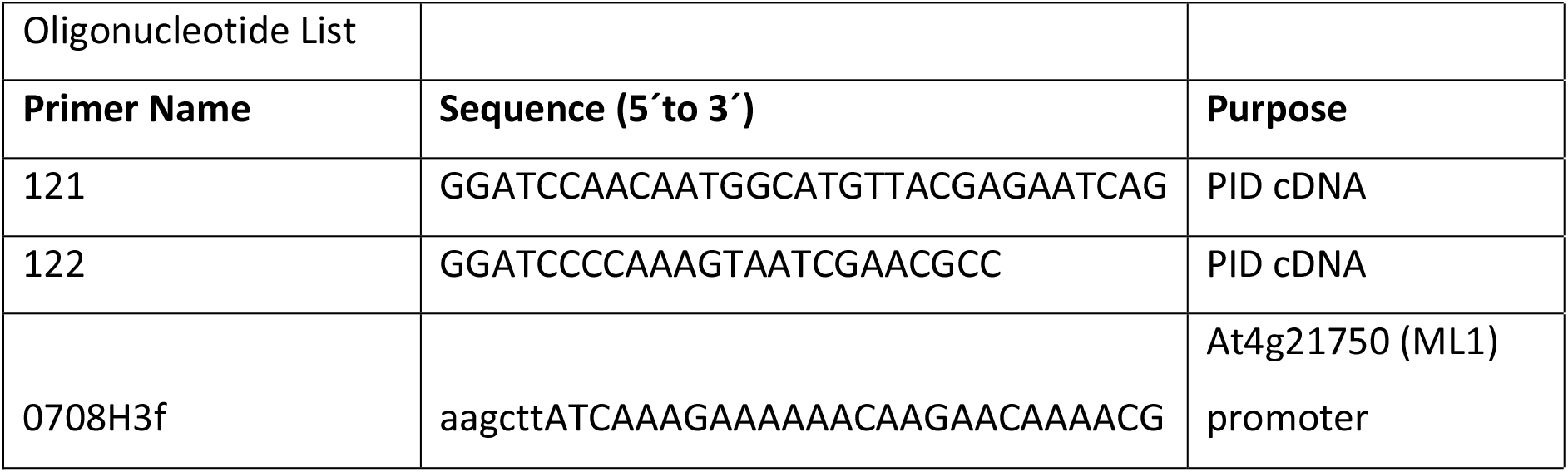
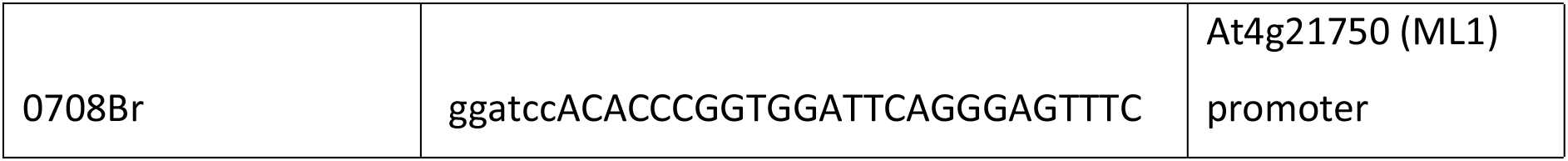

### NPA and DEX treatments

For NPA treatment, *pid-4 PIN1-GFP* seedlings were grown on GM agar plate until pin-like inflorescence meristem formation. The entire seedlings were then transferred onto the GM agar plate supplemented with 10μM NPA. Plants were then grown continuously on NPA containing medium and imaged under confocal and/or brightfield microscope for 3-7 days. For mock treatment, the plants were shifted from GM agar plate to GM agar with equivalent volume of DMSO.

To induce PID-2XVENUS sectors in epidermis, *pid* mutants harbouring *ML1p::CRE-GR+ UBQ10p::lox spacer lox::PID-2XVENUS* were grown on GM agar plate until the emergence of flowerless dome. Then 10-20μl of 10μM DEX in sterile water (10mM stock, dissolved in absolute ethanol) were directly applied on the dome meristem and were imaged at different time interval (till 24 hours or 96 hours). For every experiment, *pid* mutants were imaged before and after induction.

### Sample preparation for confocal live imaging

For Fig. 2a, wild-type inflorescence meristem was dissected and mounted on GM agar plate as previously described (Heisler and Ohno, 2014). For *pid* mutant meristem imaging, the leaves of the mutant plant covering the dome meristem were dissected away under water and then mounted the whole plant with roots on GM agar plate. Prior imaging, the mounted plants were kept submerged under sterile water for 30 minutes to halt meristem growth. For high resolution images (for Fig. 2k-m), meristem was briefly fixed in 4% paraformaldehyde as described previously(Heisler et al., 2005a) and then plasmolysed in 1M Sucrose for 1h. The plasmolysed meristem was imaged after mounting the meristem on GM agar plate filled with sucrose solution.

### Settings for confocal live imaging and time-lapse imaging

Confocal live imaging was performed on a Leica TCS-SP5 upright laser scanning confocal microscope with hybrid detectors (HyDs) using a 25X water objective (N.A 0.95) or 63X objective (N.A 1.20). Either 512×512 or 1024×1024 (for high resolution images) pixel format was used. Bidirectional scan was set with a scan speed of 400Hz or 200Hz. Line averaging used was 2 or 3. The thickness of the optical sections was 1μm.

Argon laser was used for both GFP and VENUS. GFP was excited using 488nm laser and emission window was set to 493-512nm. VENUS was excited with 514 nm laser and detected using a 520-560 nm window. Pinhole was adjusted depending on the fluorescence brightness to prevent signal saturation or bleaching. Smart gain was set to 100%. Sequential scan mode with switching in between-frames was used for imaging GFP and VENUS together.

To monitor the shift in PIN1 polarities with respect to PID clones, a time lapse imaging experiment was performed on *pid* mutant harbouring *ML1p::CRE-GR+ UBQ10p::lox spacer lox::PID-2XVENUS*. The time lapse imaging was followed at every 1 hour or 2 hours interval for a duration of 12 hours or 24 hours. PIN1-GFP expression was monitored at all time intervals, but PID-2XVENUS expression was monitored only at 2 or 3 intervals to minimize photo-toxicity. After every scan, the plants were quickly placed back under light after removing the water.

### Image analysis and data processing

The images were analysed using Imaris 9.1.2 (bit-plane), or Image J (FIJI, https://fiji.sc). The images were annotated and arranged in Adobe Photoshop 2020. PIN1 polarity assessments were made according to the presence of arcs of GFP signal extending beyond cell junctions and around cell corners.

## Acknowledgements

This research was supported by the Australian Government through the Australian Research Council’s Discovery Projects funding scheme (project DP180101149) awarded to MGH. Preliminary work was supported by the European Molecular Biology Laboratory, Heidelberg. We would like to thank Mary Byrne for helpful discussions.

## Author Contributions

AK performed most experiments and helped write the manuscript. NB performed some experiments and provided materials. CO provided materials and helped write the manuscript. MGH conceived the research and helped write the manuscript.

---

**Figure S1.**
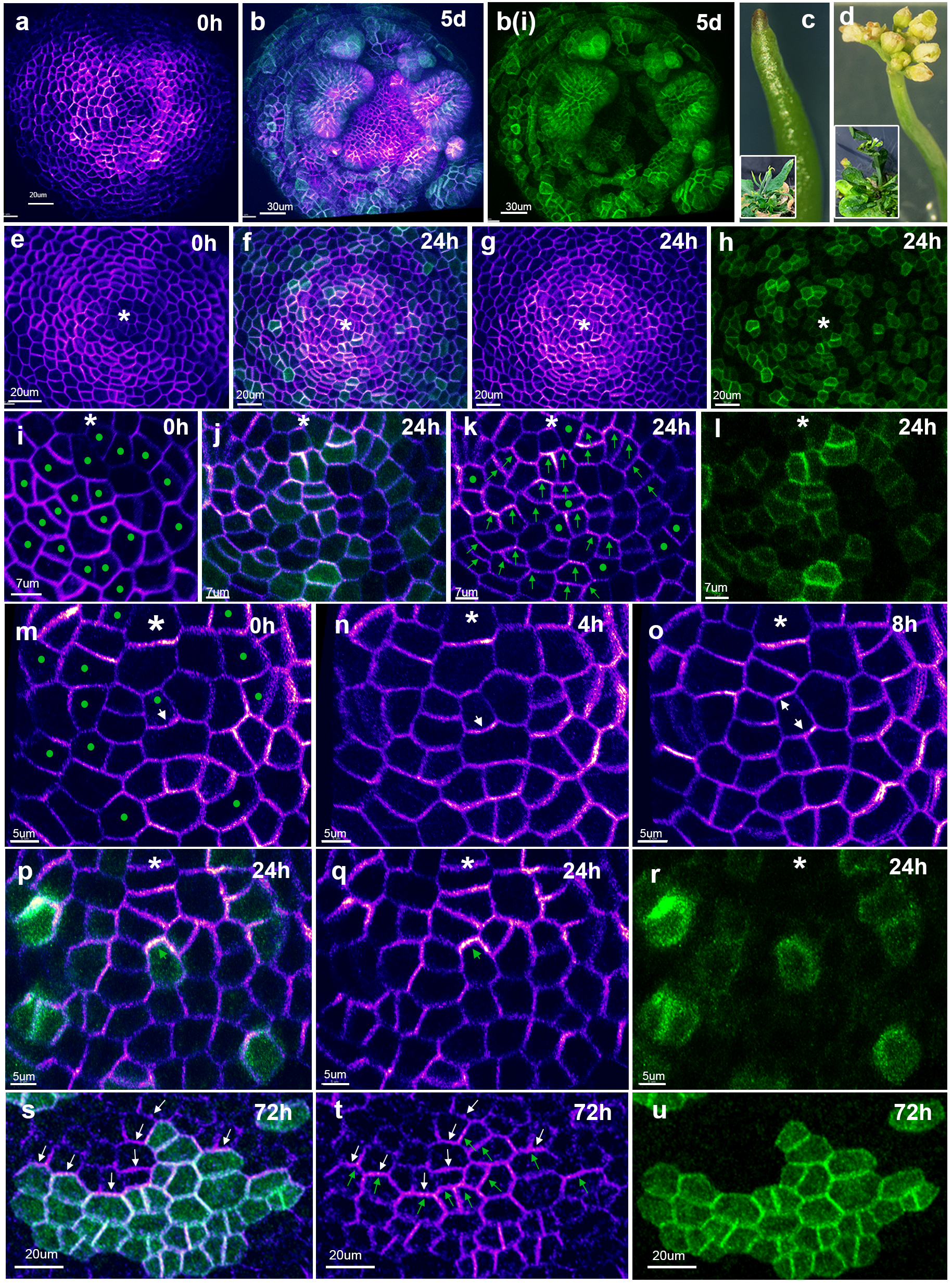
PID promotes changes in PIN1 polarity irrespective of initial or neighboring cell polarities. **a**, *pid-4* meristem expressing PIN1-GFP (magenta) before induction of PID-2V. **b**, Same *pid-4* meristem expressing PIN1-GFP (magenta) as in (**a**) 5 days after clonal induction of PID-2V (green). Note developing organs marked by large sectors of PID-2V expression. **b(i)**, Same as (**b**) showing PID-2V (green) alone. **c,** pin-like structure of uninduced *pid-4 PIN1-GFP* inflorescence. **d,** Inflorescence of *pid-4 PIN1-GFP* rescued by clonal expression of PID-2V after 1 week. **e-r,** Images of the *pid-4* meristem before (**e,i,m**) and after (**f-h, j-l, n-r**) induction of *PID-2V* (green) expressing clones showing changes in *PIN1-GFP* (magenta) polarity. **i-l,** Magnified images of the same meristem in (**e-h**). **m-r**, Magnified high resolution time-series images of the same meristem from **Figure 2** (main text) **e-j**. Note apical shift in PIN1-GFP polarity (green arrows in **k,p,** and **q**) in cells expressing PID-2V (green dots in **i,m**). Green dots in (**k**) mark cells with undiscernible polarity. Note a gradual shift in PIN1-GFP (magenta) polarity from basal (**m,n**) to bipolar (**o**) to apical (**p,q**) over time after induction of PID-2V (green). Arrows indicate observed polarity, i.e., downward= basal, upward= apical, two arrows in opposite directions = bipolarity. Asterisk marks middle of the meristem. **s-u**, Flank of a *pid-4* shoot apex showing opposing PIN1-GFP (magenta) polarities (green and white arrows in (**s**) and (**t**)) in adjacent cells due to differential PID-2V expression (green) (**s,u**) 3days after induction.

**Movie1:** Volume rendering of primordium depicted in **Figure 2 b-d** with different channel combinations displayed over time.

**Movie 2:** Animated series of four time points depicted in **Figure 2 e-j** showing shift in PIN1-GFP (magenta) polarity from basal to apical in PID-2V (green) expressing cells in *pid-4* mutant meristem.

